# GRIP: physics-informed neural network for gradient retention time prediction in liquid chromatography

**DOI:** 10.1101/2024.11.11.622855

**Authors:** Kevin George, F.P. Jake Haeckl, Gerrit Großmann, Alexey Gurevich, Azat Tagirdzhanov

## Abstract

Gradient liquid chromatography has numerous applications in life sciences. Retention time prediction remains challenging due to complex underlying physical processes and highly variable chromatographic conditions. Here, we present GRIP, a physics-informed neural network for gradient retention time prediction that explicitly uses experimental setup parameters. GRIP demonstrates zero-shot generalization to unseen chromatographic systems while being on par or out-performing the transfer learning-based baseline. This approach can computationally guide the experimental setup configuration tailored to specific compounds of interest.

## 1. Introduction

Gradient high-performance liquid chromatography (HPLC) is a widely used analytical technique for separating compounds in a complex chemical mixture. Often coupled with mass spectrometry, it became a method of choice in various areas of life science, including metabolomics (Harrieder et al., 2022), environmental science (Escher et al., 2020), pharmaceutical (Chew et al., 2021) and medical research (Wu & French, 2013), as it enables the identification of small molecules in complex samples by comparing them to reference standards.

During separation, the sample is passed through a chromatographic column and separated in time depending on the interaction of its component compounds with the column. The time at which the compound leaves the column is called retention time (RT). The experimental derivation of RT is costly and requires efforts to isolate and purify compounds. In-silico prediction of the RT from molecular structure enables the use of chemical databases. This allows to broaden the range of compounds that can be identified and will assist in-silico identification of compounds in mass spectrometry data, providing additional information that helps improve the accuracy of molecular identification (Nothias et al., 2020; Mullowney et al., 2023).

A compound’s retention time depends on the HPLC system parameters, such as column properties, gradient program, and temperature, which makes its prediction extremely difficult. In recent years, several approaches using traditional machine learning and deep learning have emerged to predict RT. For instance, Barron & McEneff (2016) used molecular descriptors and an artificial neural network to predict RT. Bonini et al. (2020) developed Retip, which used six different machine learning models for predicting RT. However, these methods are limited to a particular HPLC system that was used to generate their training data, making them system-dependent. To apply these models to a different HPLC system, a sufficient amount of new system-specific data is required, followed by retraining the model on this data.

In contrast to system-specific approaches, Stanstrup et al. (2015) used RT data from multiple chromatographic systems to develop a projection model that maps RT from one system to another. Building on this Bouwmeester et al. (2020) developed CALLC, which combines machine learning with a projection model (termed *calibration curve*) to predict RT across different systems. Another approach, one-hot encoding of various setups alongside molecular descriptors to train a multi-layer perceptron for RT prediction (Pasin et al., 2021).

In 2019, Domingo-Almenara et al. introduced SMRT, a large-scale dataset that covers 80,038 small molecules and trained a deep learning model for RT prediction. Following the release of the SMRT dataset, numerous studies emerged, exploring deep learning methods for RT prediction. In one such study the authors proposed GNN-RT, a graph neural network that predicts RT directly from molecular structure (Yang et al., 2021). Fedorova et al. (2022) used 1D-CNNs on SMILES strings and Xue et al. (2024) introduced RT- Transformer, which combines graph attention networks with 1D-Transformers to learn molecular representations. More recently, Vik et al. (2024) evaluated different combinations of molecular fingerprints and descriptors using models such as XGBoost (Chen & Guestrin, 2016), AttentiveFP (Xiong et al., 2019), and ChemProp (Heid et al., 2023), concluding that ChemProp with RDKit descriptors (Landrum, 2016) performed the best. These deep learning methods leverage pre-training on the SMRT dataset, followed by fine-tuning on system-specific datasets, to demonstrate their transferability across different HPLC systems. In contrast to the transfer-learning methods, García et al. (2022) adopted a projection method alongside a deep learning model trained on the SMRT dataset to predict RT across various systems. However, both the fine-tuning and projection-based approaches cannot predict RT on a new or reconfigured HPLC system without a set of additional calibration measurements.

In this work, we employ the physical principles of liquid chromatography to create GRIP, a deep neural model that explicitly uses HPLC system parameters to predict retention time. We train the model on 65 reverse-phase HPLC datasets from the RepoRT repository (Kretschmer et al., 2024). Our results show that GRIP can generalize to unseen chromatographic systems without any fine-tuning. In our benchmark, GRIP achieved better or comparable results with a transfer learning-based method, GNN-RT (Yang et al., 2021), used as a baseline.

## 2. Theoretical background

The chromatographic column is packed with adsorbent particles, called the stationary phase. The sample of interest (analyte compounds) is mixed with a solvent (commonly referred to as mobile phase), and then travels through the column. Each analyte in the mixture interacts differently with stationary and mobile phases, resulting in different retention times. The mobile phase is often a mixture of organic solvents and water (Snyder et al., 2009).

In the isocratic HPLC, when the fraction *φ* of the organic solvent in the mobile phase is constant, the retention time of the analyte can be expressed as

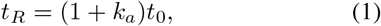

where *t*_0_ is the holdup time of the column, and *k*_*a*_ is the retention factor. The retention factor depends on the analytes’ properties and the system parameters, such as temperature and the mobile phase composition *φ* (den Uijl et al., 2021).

The holdup time *t*_0_ is the time it takes for the mobile phase to pass through the column. Even though this parameter is important for modeling, it is rarely reported in experimental papers. It can be expressed as

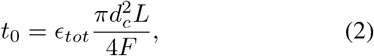

where *d*_*c*_ and *L* are the internal diameter and the length of the column, *F* is the flow rate, and *ϵ*_*tot*_ is the total porosity of the column (Snyder et al., 2009). In practice, *ϵ*_*tot*_ is usually calculated from the retention time of an unretained molecule that has no affinity to the stationary phase of the column (Rimmer et al., 2002).

In gradient HPLC the mobile phase composition changes over time according to the gradient program. In this case, we can get *t*_*R*_ from the fundamental equation for gradient elution (Nikitas & Pappa-Louisi, 2005),

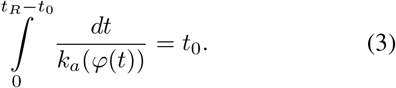

Here, *k*_*a*_(*φ*) is the retention factor in the isocratic mode corresponding to the constant mobile phase composition equal to *φ*. The gradient program *φ*(*t*) is usually assumed to be a linear function of time (Nikitas & Pappa-Louisi, 2005; den Uijl et al., 2021).

To utilize (3) for the retention time prediction, it is crucial to model *k*_*a*_(*φ*). Over the years, several empirical models emerged, that were developed having two objectives in mind: first, they have to be powerful enough to fit the data, and, second, they make the integral in (3) analytically tractable. For a review on the modelling efforts, please refer to (den Uijl et al., 2021). In our approach, we rely on the linear solvent strength (LSS) model, which has the form

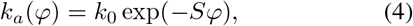

where *k*_0_ and *S* are empirical coefficients specific for a particular analyte and an HPLC system setup (Snyder et al., 1979).

## 3. Methods

### 3.1. Piecewise linear gradients

In practice, gradient programs often consist of multiple linear segments (Snyder et al., 2009),

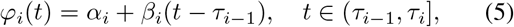

where *τ*_*i*_, *i* = 1, …, *n, τ*_0_ = 0, are the time steps of the gradient, and *β*_*i*_ is the slope. In this case, substituting (4) and (5) into (3), we can rewrite it as

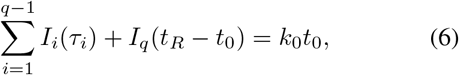

where *I*_*i*_ is the integral over the *i*-th segment of *φ*(*t*), and *q* is the index of the segment during which the analyte leaves the column (Figure 1). Changing the variable in the integrals, we immediately get

**Figure 1:**
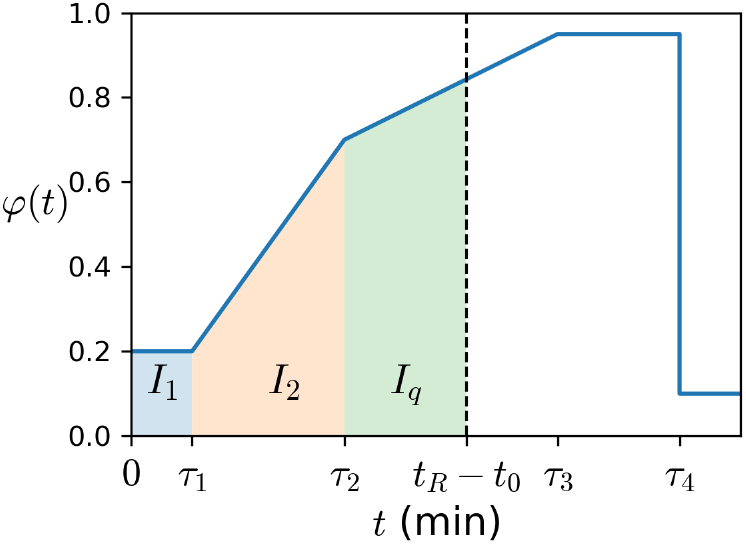
A schematic example of a gradient program. The dashed line shows *t* = *t*_*R*_ *t*_0_, highlighted areas under the curve correspond to the integrals *I*_*i*_.

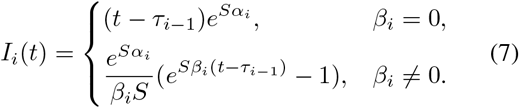

Retention time is expressed from (6) as

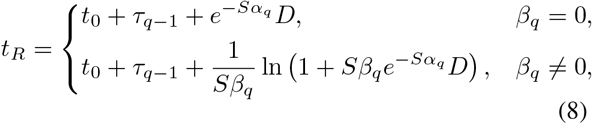

Where

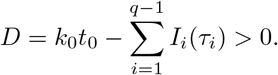

### 3.2. Model architecture

The model architecture is shown in Figure 2. It takes as input datapoints, each representing a measurement of a specific molecule within a particular HPLC system and structured as (*G*, **d, s**, *g, t*_0_, *t*_*R*_):

**Figure 2:**
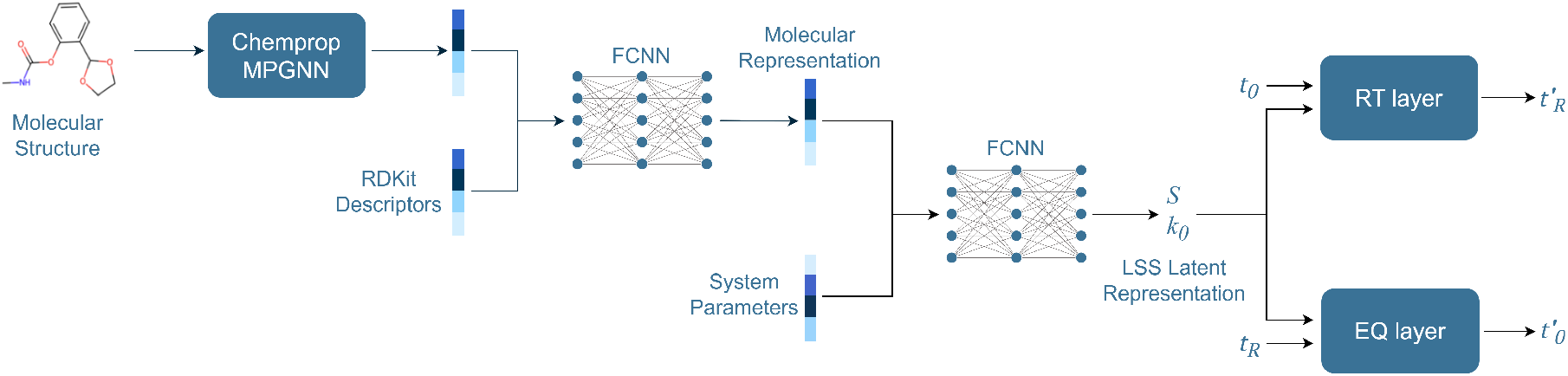
Model Architecture. Molecular graph is a graph representation of a molecule with atoms as nodes and bonds as edges. Molecular descriptors are computationally generated numerical representations of physical and chemical properties of the molecule, such as LogP and topological polar surface area. System parameters are the parameters specific to the experimental HPLC system, such as temperature or stationary phase of the chromatographic column.

- *G* represents the molecule, provided in the form of a molecular graph,
- **d** is a vector of molecular descriptors generated with RDKit (Landrum, 2016),
- **s** is a vector of system parameters,
- *g* represents the gradient program (5),
- *t*_0_ is the holdup time, computed from the column parameters using (2),
- *t*_*R*_ is the retention time, which serves as the target variable for prediction.

First, we use a message passing graph neural network (MPGNN) architecture from Chemprop (Heid et al., 2023) to learn the embedding vectors of the molecular graphs *G*. MPGNN output

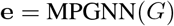

is concatenated with vectors of molecular descriptors **d** and passed through fully connected layers to obtain the final molecular representation

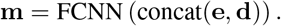

The molecular representation is concatenated with a vector of system parameters **s**,

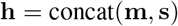

and the LSS model parameters are predicted with a fully-connected network,

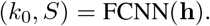

The parameters and *t*_0_ are then passed to the layer that uses (8) for producing the final prediction,

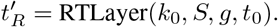

In addition, during training the left-hand side of the equation (6) is computed given the ground truth value of *t*_*R*_,

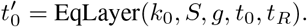

During training, the value of the gradient segment *q* is determined from the ground truth value of *t*_*R*_,

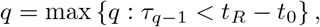

as it improves convergence. During inference, *q* is determined as

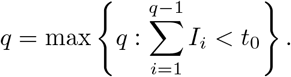

We train the model in an end-to-end manner by minimizing the loss function

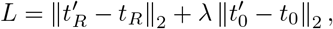

where *λ* is a hyperparameter.

### 3.3. Datasets

RepoRT is a repository of retention time datasets, measured on different instruments and in various chromatographic conditions (Kretschmer et al., 2024). It provides standardized metadata on the chromatographic system parameters as well as molecular structure of the analytes.

#### Dataset preparation

For training and evaluation of GRIP, we select reverse-phase HPLC datasets with a constant flow rate and water-acetonitrile mobile phase, since this setup is the most represented among RP datasets in RepoRT (Kretschmer et al., 2024). To account for different stationary phases, we enrich the metadata with the hydrophobic-subtraction model (HSM) parameters based on the manufacturer and model of the HPLC column used to generate the dataset (Snyder et al., 2004). As the HSM parameters for one of the columns, Restec Raptor Biphenyl, are missing, we use Kinetex Biphenyl as a substitute, as it is listed by the manufacturer among similar stationary phases (Res, 2022). We discard datasets with incomplete metadata and stationary phases covered by less than 200 datapoints.

#### Dataset composition

The resulting dataset consists of 74 RepoRT datasets obtained from 6 different data sources according to the RepoRT metadata, and generated using 30 experimental setups with 8 stationary phases. Dataset details are listed in Supplementary Table S1. The dataset comprises 44,973 retention time measurements for 11,903 unique molecules. The distribution of chemical classes is shown in Supplementary Figure S2. The column temperature ranges from 30 to 50^*°*^C. Gradient programs are shown in Supplementary Figure S1 and range from 8 to 20 minutes. Pairwise comparisons between stationary phases computed with the column selectivity function (Snyder et al., 2004) are shown in Supplementary Figure S3.

#### SMRT dataset

The SMRT dataset (Domingo-Almenara et al., 2019) is a large-scale dataset covering 80,038 small molecules, available in RepoRT under ID 0186. We solely use this dataset to pretrain the baseline method, GNN-RT (Yang et al., 2021), which is based on transfer learning. We do not include this dataset into GRIP’s training data, as it does not provide all the required system parameters.

### 3.4. Generating synthetic data

We generate additional synthetic datapoints to improve regularization during training. The generating procedure relies on mapping retention time measurements between datasets using calibration curves first introduced in (Stanstrup et al., 2015).

For the target datasets in our training data, we selected source datasets from RepoRT with at least 50 molecules in common and fitted a generalized additive model as described in Figure 3 and Supplementary Note S1. We selected the regions of applicability based on the number of datapoints, confidence interval of the model and the number of outliers. We kept only the datapoints supported by more than one source dataset with a standard deviation of the predicted value of less than 1 minute. The procedure is described in detail in Supplementary Note S1.

**Figure 3:**
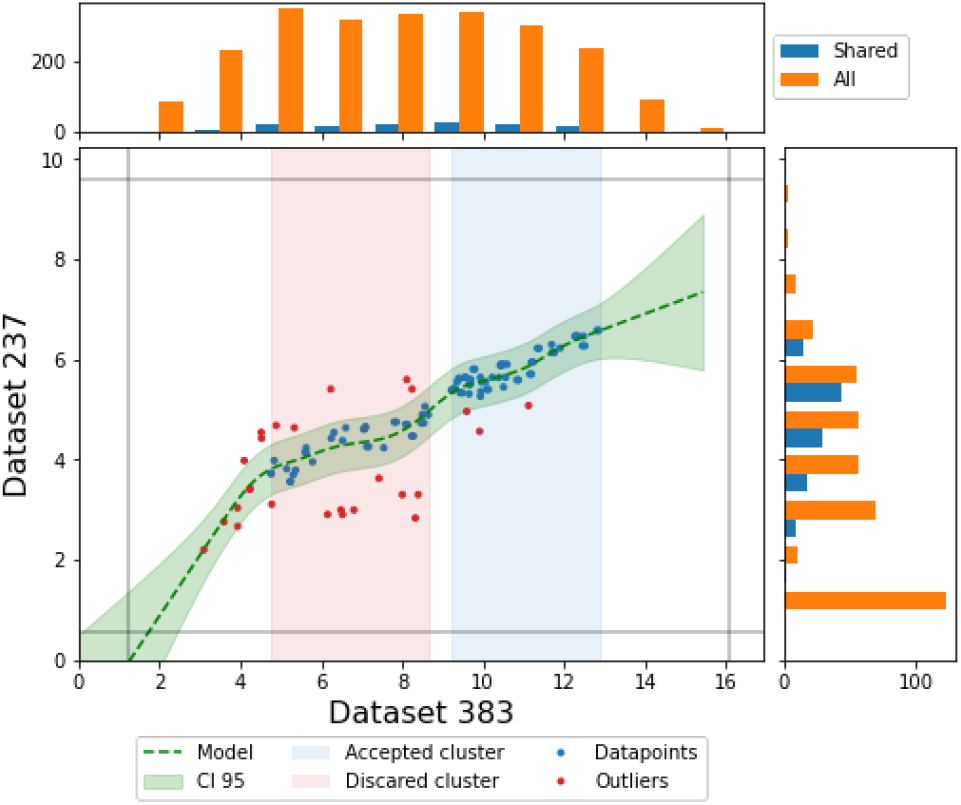
A calibration curve mapping RT from Dataset_383_ to Dataset_237_ (dashed line). Dots represent molecules shared between the two datasets. Light green area shows the 95% confidence interval computed from the model. The blue shaded area shows the region meeting all applicability criteria, while the red shaded area shows the region that is supported by enough datapoints, but is still discarded due to a high number of outliers. Histograms show distribution of RT in each of the datasets.

### 3.5. Experimental setup

Our experiments aim to estimate the generalization performance of the model (i) across chemical space, and (ii) across different chromatographic conditions.

We report the results of two key experiments:

- **Experiment 1: Generalization to New Molecules** – This experiment tests the model’s ability to generalize to novel molecules that differ significantly from those encountered during training.
- **Experiment 2: Generalization to New System Parameters** – This experiment evaluates the model’s generalization capacity to previously unseen system parameters. In particular, we assess the model’s zero-shot generalization capability for new experimental setups and molecules both similar and not similar to those encountered in the training phase.

We now explain how the dataset is partitioned for Experiments 1 and 2. The whole process is illustrated in Figure 4.

**Figure 4:**
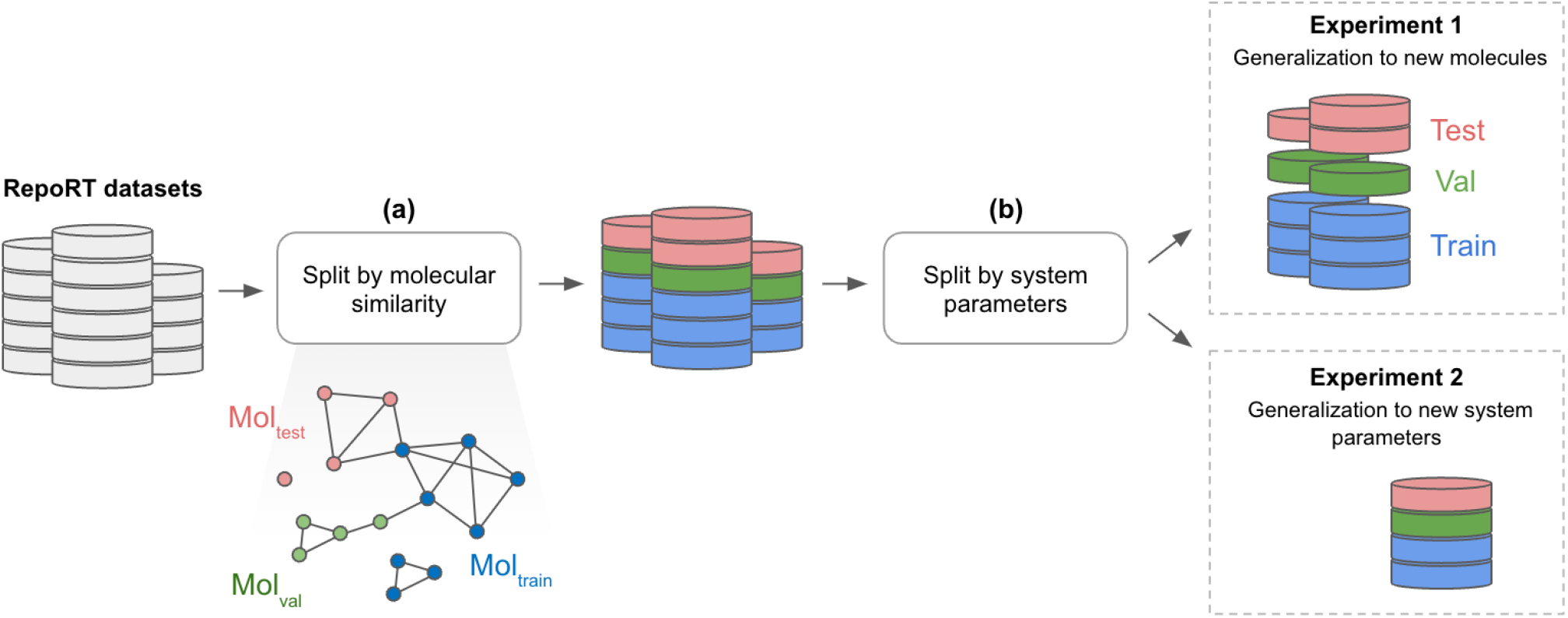
Data partitioning process. **a** The first splitting uses hierarchical clustering of molecules based on Tanimoto similarity, dividing each RepoRT dataset into three splits of significantly different molecules. The datasets are divided in a coordinated way, forming three disjoint sets of molecules, Mol_train_, Mol_val_, and Mol_test_, which aggregate molecules from combined training, validation, and testing splits, respectively. **b** The second splitting is based on HPLC system parameters. This approach enables model performance evaluation under different scenarios: seen systems with unseen molecules (Experiment 1); unseen system with seen molecules, and unseen system with unseen molecules (Experiment 2).

#### Splitting by molecular similarity

Our goal is to maintain the same split of molecules in the GRIP training data, SMRT dataset, and other RepoRT datasets not included in the GRIP training data but used for generating synthetic datapoints. We start with a set of all unique molecules in the combined RepoRT datasets and split it into the subsets Mol_train_, Mol_val_, and Mol_test_ based on molecular similarity.

Specifically, we compute pairwise similarities between the molecules (Tanimoto similarity computed from Morgan fingerprints, radius 3) and create a similarity network with nodes connected by a weighted edge if their similarity is greater than 0.5. We group the molecules by clustering network nodes with a community detection algorithm (Raghavan et al., 2007). Each datapoint in RepoRT datasets is then assigned a cluster ID. We split all the data to 80:10:10 ratio in a stratified group manner using cluster IDs as groups and RepoRT IDs as strata. This approach ensures that all datapoints associated with one group of molecules are assigned to the same split, and individual RepoRT datasets are evenly represented across the splits.

#### Splitting by system parameters

In this step, we create new test datasets specifically for Experiment 2, which evaluates generalization capabilities with respect to new system parameters. To achieve this, we identify two specific system parameters and remove all associated datapoints from the training, validation, and test sets, placing them instead into the following evaluation datasets:

- Dataset_310*−*317_ is formed of 8 RepoRT datasets generated with the Waters CORTECS C18 column. These datasets differ one from another only by temperature and the set of analyzed molecules. At the same time, the gradient program used to generate this dataset is present in training data.
- Dataset_389_ is the only dataset coming from the “Publication - Aalizadeh” source and was generated with a gradient program dissimilar to all the datasets in training, see Supplementary Figure S1.

This results in the following dataset splits: training set (33,859 datapoints for 9,545 unique molecules), validation set (4,176 datapoints for 1,111 unique molecules), and testing set (4,193 datapoints for 1,193 unique molecules). We extend the training set by adding synthetic datapoints generated as described in Section 3.4. To avoid data leakage, we only add synthetic datapoints for the molecules in the Mol_train_ subset. This contributes 72,472 datapoints for 4,274 unique molecules, of which 984 were not present in the original training set. Training, validation, and testing sets comprise datapoints from 65 RepoRT datasets, while 9 RepoRT datasets are hold out for evaluation in the Experiment 2.

We train the model in an end-to-end manner using column temperature and HSM parameters of the stationary phase as a set of system parameters. The validation set is used to determine the model hyperparameters. We use the model that achieved the best results on the validation set to evaluate its performance on both the testing set (Experiment 1) and evaluation datasets (Experiment 2).

## 4. Results

### 4.1. Generalization to new molecules

Here we report the performance of the model on the testing set consisting of the molecules not seen in training, with a mean similarity to the closest molecule in the training set of 0.4. The distribution of molecular similarities is shown in Supplementary Figure S4.

Table 1 summarizes the performance of the model in terms of mean and median absolute errors (MAE, MedAE), mean and median relative errors (MRE, MedRE), and the coefficient of determination *R*^2^. The results for individual RepoRT datasets are listed in Supplementary Table S2. We exclude predictions on synthetic datapoints when computing performance metrics. On the testing set GRIP shows an

**Table 1:** Retention time prediction accuracy on training, validation, and testing sets.

|  | Training | Validation | Testing |
| --- | --- | --- | --- |
| # Datapoints | 33,859 | 4,176 | 4,193 |
| # Molecules | 9,545 | 1,111 | 1,193 |
| MAE (min) | 0.25 | 0.41 | 0.42 |
| MRE | 0.10 | 0.16 | 0.16 |
| MedAE (min) | 0.14 | 0.24 | 0.26 |
| MedRE | 0.06 | 0.10 | 0.12 |
| $R^2$ | 0.96 | 0.87 | 0.88 |

MAE of 0.42 min and *R*^2^ of 0.88. This suggests that the model was able to generalize well to dissimilar molecules. We show the performance of the model on the training set as a reference for the next section.

### 4.2. Generalization to new system parameters

We assess zero-shot generalization capability of GRIP by evaluating it on the datasets having HPLC system parameters not seen in training. The results are reported in Table 2.

**Table 2:** Retention time prediction accuracy on evaluation datasets. The second row shows the number of unique molecules and the overlap with the molecules seen in training (in parentheses). The results are reported separately on two subsets: Mol_train_, containing molecules similar to those seen in training, and Mol_test_, consisting of dissimilar molecules.

|  | Dataset<sub>389</sub> |  | Dataset<sub>310–317</sub> |  |
| --- | --- | --- | --- | --- |
| | $\text{Mol}_{\text{train}}$ | $\text{Mol}_{\text{test}}$ | $\text{Mol}_{\text{train}}$ | $\text{Mol}_{\text{test}}$ |
| # Datapoints | 65 | 9 | 2,097 | 287 |
| # Molecules | 65 (30) | 9 (0) | 548 (546) | 70 (0) |
| MAE (min) | 1.08 | 1.43 | 0.25 | 0.64 |
| MRE | 0.30 | 0.15 | 0.11 | 0.23 |
| MedAE (min) | 0.79 | 1.28 | 0.20 | 0.39 |
| MedRE | 0.11 | 0.10 | 0.07 | 0.21 |
| $R^2$ | 0.92 | 0.87 | 0.96 | 0.78 |

Dataset_310*−*317_ was generated using a stationary phase different from those seen in training and a seen gradient program. As seen from Table 2, the performance of GRIP on the Mol_train_ subset is similar to its performance on the training data. On Mol_test_ the model shows a higher MAE of 0.64 min and a lower *R*^2^ of 0.78.

Dataset_389_ was generated with a gradient program dissimilar gradient programs seen in training. GRIP demonstrates higher MAE of 1.08 min on the Mol_train_ subset and MAE of 1.43 min on the Mol_test_ subset, while the coefficient of determination remains comparable to the training results. Still, GRIP achieves acceptable results.

### 4.3. Comparison with transfer learning approach

We benchmark our model against GNN-RT (Yang et al., 2021), a transfer learning method that relies on a similar GNN encoder to learn molecular representations. Following (Yang et al., 2021), we pre-trained GNN-RT on the SMRT dataset (Domingo-Almenara et al., 2019) using the source code from the original publication. For a fair comparison with GRIP, we followed the same split of the chemical space in the SMRT dataset as in our training data, in contrast to the random split used in (Yang et al., 2021). GNN-RT achieved on the testing subset of SMRT a MAE of 0.91 min.

We fine-tuned GNN-RT on each of the evaluation datasets using datapoints associated with the molecules from the Mol_train_ subset. To model a real-world scenario, where the number of measurements for an HPLC system of interest is often limited, we measured the performance of GNN-RT depending on the amount of data available for transfer learning. For each dataset size, we randomly sampled datapoints from the Mol_train_ subset for fine-tuning and tested the model on the Mol_test_ subset.

Figure 5 shows performance of GNN-RT compared to GRIP on three selected evaluation datasets. The results on the remaining evaluation datasets are shown in Supplementary Figure S5. As expected, GNN-RT performance is sensitive to the amount of data available for fine-tuning. Interestingly, GRIP outperforms GNN-RT or shows comparable performance in all the experiments without any fine-tuning.

**Figure 5:**
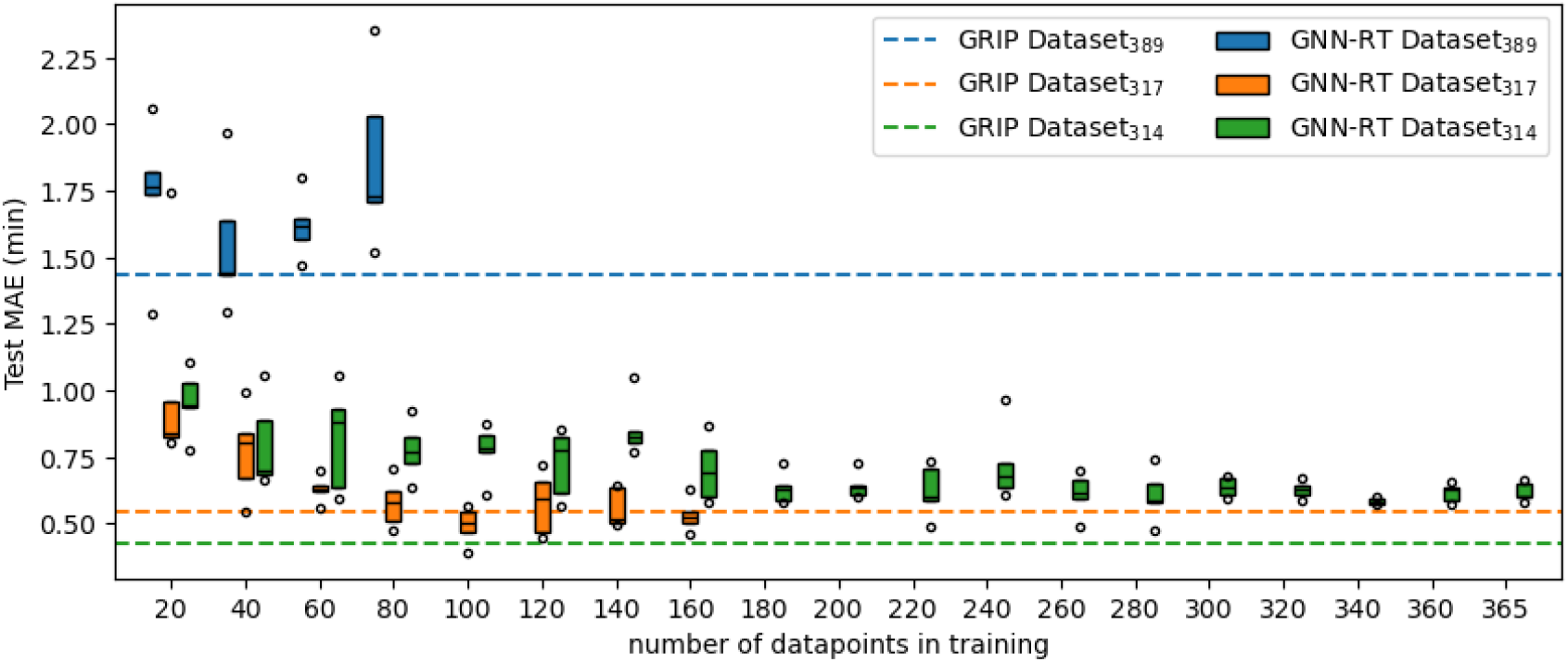
Comparison of the performance of GNN-RT (Yang et al., 2021) and GRIP (our model) across three evaluation datasets. GNN-RT was fine-tuned on varying dataset sizes, increasing by 20 datapoints for each evaluation dataset. For each dataset size, data points were randomly sampled from the Mol_train_ subset and the model was tested on the Mol_test_ subset. Box plots show the MAEs of GNN-RT on the testing subset for five repetitions of random sampling. In contrast, GRIP was directly evaluated on the Mol_test_ subsets without fine-tuning (dashed line).

## 5. Discussion

GRIP introduces a novel physics-informed deep model for retention time prediction. The model is based on the fundamental equation for gradient elution and the linear solvent strength (LSS) model and explicitly uses parameters of the experimental high-performance liquid chromatography (HPLC) system and molecular structure of a compound.

The LSS model describes the analyte retention factor as a function of solvent concentration, characterized by two parameters. While GRIP predicts the parameters of the LSS model, it is important to note that LSS parameters predicted from gradient data may not reliably match those obtained from the measurements under isocratic conditions (Ford & Ko, 1996).

We trained GRIP using a diverse selection of reverse-phase HPLC datasets featuring different chromatographic conditions with water-acetonitrile mobile phase. However, publicly available data limits our ability to evaluate the generalization of the model to unseen HPLC systems more extensively. This should be taken into account in practical applications of the model on out-of-domain data. We plan to broaden the scope of our experiments as well as benchmark GRIP against other retention time prediction models in later revisions of the paper.

In our model, we assume that the effects of the dwell volume and extra-column volume are negligible or accounted for by the initial isocratic hold in the gradient, as discussed by Dolan (2006) and Hong & McConville (2018). The RepoRT metadata does not provide neither these parameters nor the HPLC instrument. We plan to revisit this aspect in the future.

Our experiments show that the model is better or at par with the baseline transfer learning method without requiring additional data. We plan to cover a broader range of HPLC methods and test its performance on more diverse setups. However, GRIP can already assist with identifying compounds in biological samples and can also be used for simulating chromatographic experiments for designing HPLC system configurations to suit the particular compounds of interest.

