## Supplementary Material for "GRIP: physics-informed neural network for gradient retention time prediction in liquid chromatography"

### Supplementary Notes

#### Supplementary Note S1 Generating synthetic data

For each target dataset we selected source datasets from RepoRT that have at least 50 molecules in common and at least 10 molecules that have not been measured in the target dataset. We fitted calibration curves between the selected pairs with a monotonically constrained generalized additive model (LinearGAM implementation from the pygam module v.0.9.1 [1]) following the weighting procedure described in [2]. To find the regions that can be used for generating new data, we clustered the source datapoints with 1D DB-SCAN (scikit-learn v.1.5.2 [3] implementation; eps=0.5, min_samples=5) and selected the clusters that are supported by at least 5 datapoints, have less than 10% datapoints with fitting error more than 0.5 minutes, and are at least 0.5 minutes long.

We used the models to map measurements from all RepoRT datasets to the datasets in our training data. We kept only the datapoints that are supported by more than one source dataset with standard deviation of the predicted value less than 1 minute. The results are summarized in Supplementary Table S1.

### Supplementary Figures

Figure S1: Gradient programs of the datasets grouped by the dataset source according to the RepoRT metadata (see Supplementary Table S1 for the list of datasets). Here, "Publication - Aalizadeh" gradient is used only for evaluation.

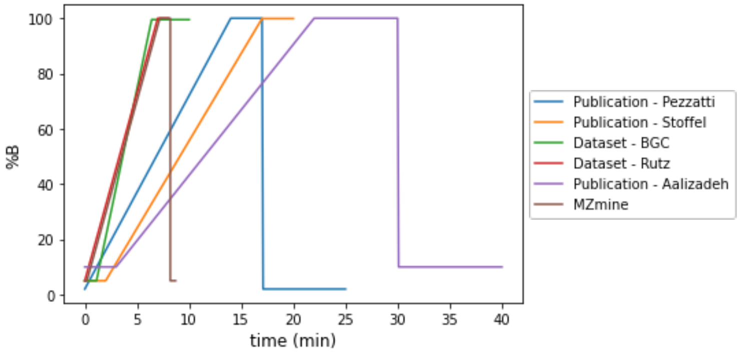


Figure S2: Distribution of ClassyFire classes across the splits. Only classes represented by at least 50 molecules are shown.

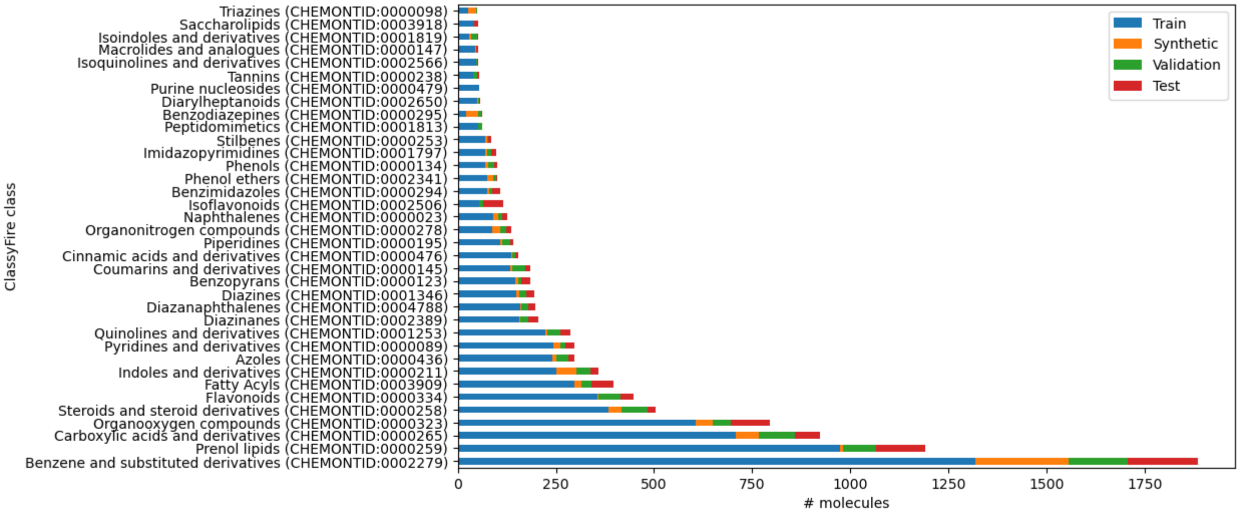

Figure S3: Similarity factors of stationary phases computed from the HSM parameters with the column selectivity function using default weighting parameters [4]. Similarity factor acts as a dissimilarity measure with the smallest values showing higher similarity between stationary phases.

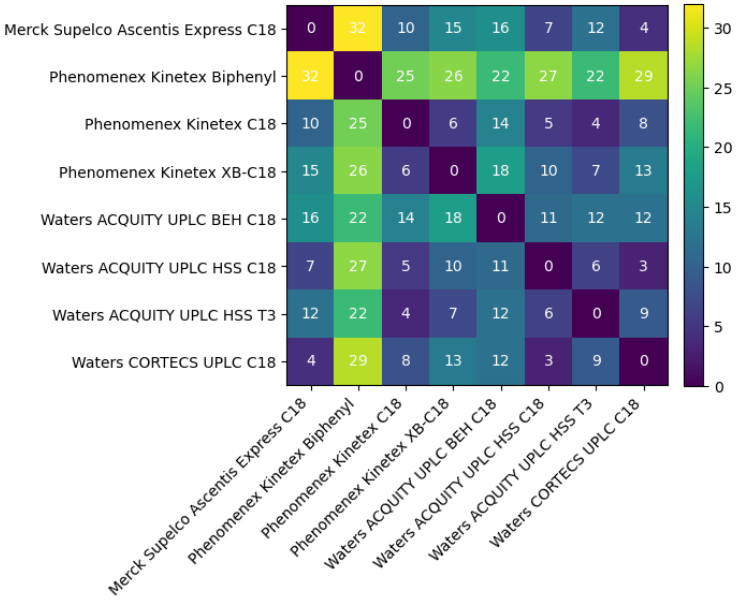


Figure S4: Distribution of Tanimoto similarities of the test and validation molecules to the closest molecule in the training set.

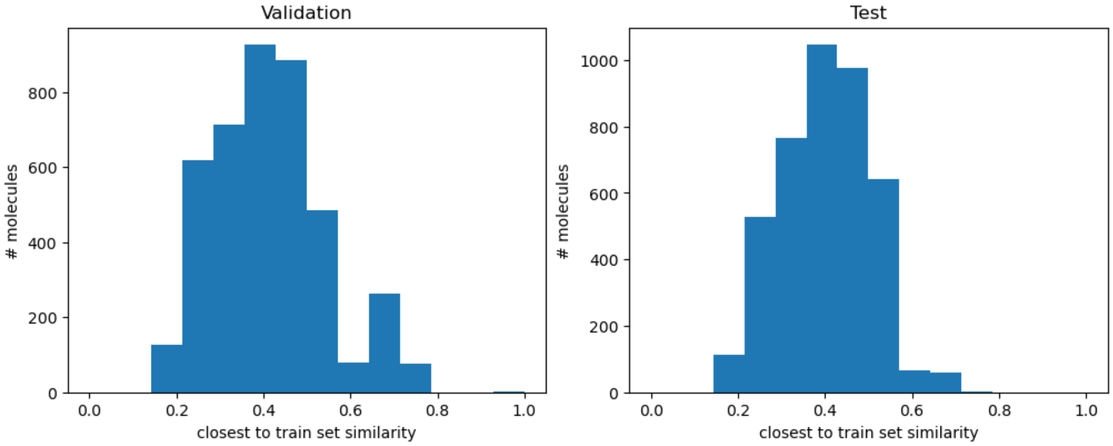

Figure S5: Performance of GNN-RT [5] and GRIP on $Mol_{test}$ subset of evaluation datasets. GNN-RT was fine-tuned on $Mol_{train}$ subsets of varying sizes, while GRIP was evaluated in zero-shot setting.

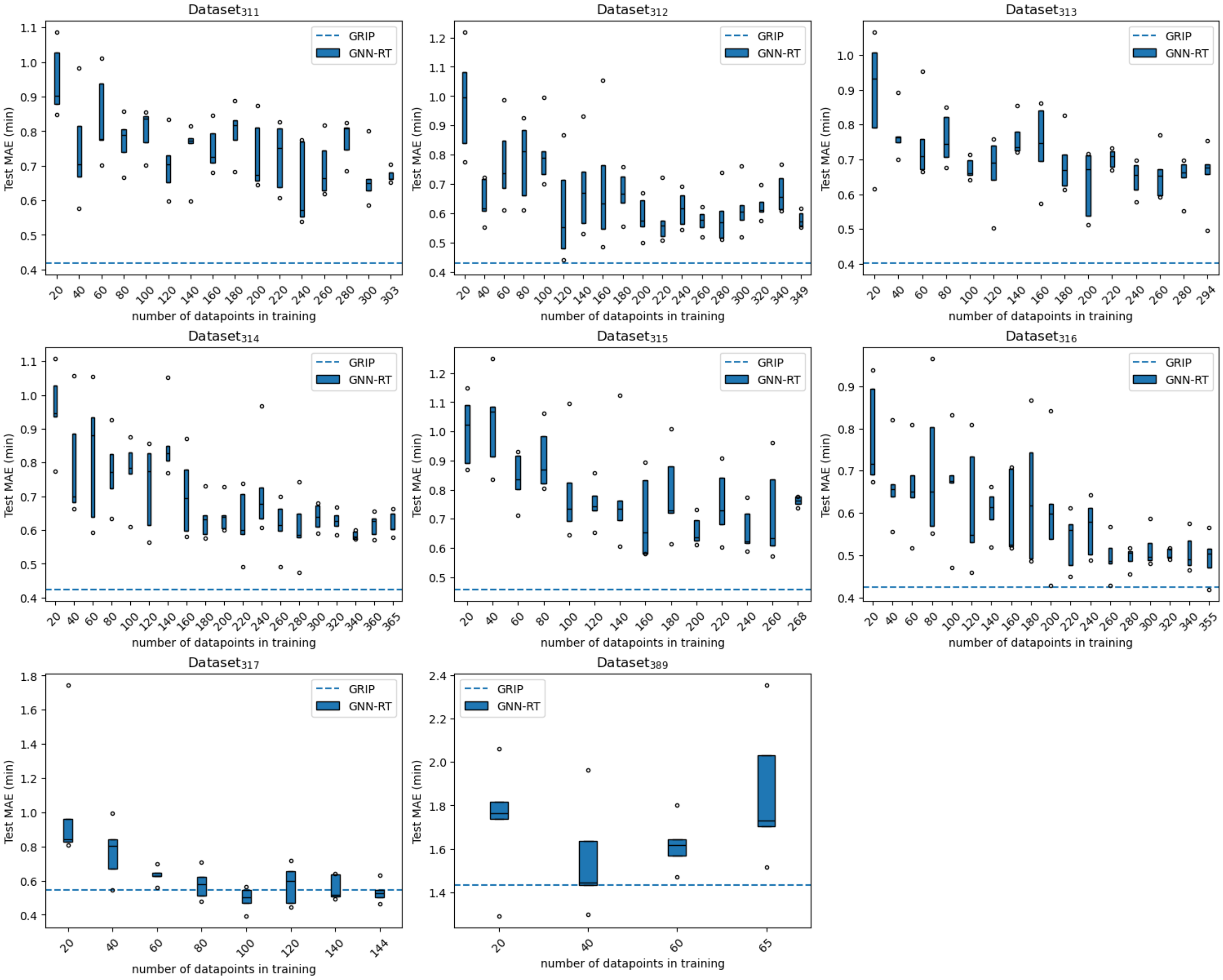

### Supplementary Tables

Table S1: Datasets used in this study

| Source | RepoRT ID | Chromatographic conditions | Number of datapoints | | | |
|---|---|---|---|---|---|---|
| | | | $Mol_{train}$ | Synthetic | $Mol_{val}$ | $Mol_{test}$ |
| **Evaluation datasets** | | | | | | |
| Publication - Aalizadeh | 0389 | Waters ACQUITY UPLC BEH C18 100x2.1mm, 1.7um, 30°C, 0.35mL/min | 65 | 0 | 9 | 9 |
| Dataset - BGC | 0310 | Waters CORTECS UPLC C18 100x2.1mm, 1.6um, 30°C, 0.4mL/min | 19 | 0 | 2 | 2 |
| | 0311 | Waters CORTECS UPLC C18 100x2.1mm, 1.6um, 30°C, 0.4mL/min | 293 | 0 | 145 | 156 |
| | 0312 | Waters CORTECS UPLC C18 100x2.1mm, 1.6um, 40°C, 0.4mL/min | 347 | 0 | 283 | 279 |
| | 0313 | Waters CORTECS UPLC C18 100x2.1mm, 1.6um, 40°C, 0.4mL/min | 285 | 0 | 153 | 164 |
| | 0314 | Waters CORTECS UPLC C18 100x2.1mm, 1.6um, 50°C, 0.4mL/min | 364 | 0 | 289 | 284 |
| | 0315 | Waters CORTECS UPLC C18 100x2.1mm, 1.6um, 50°C, 0.4mL/min | 261 | 0 | 146 | 152 |
| | 0316 | Waters CORTECS UPLC C18 100x2.1mm, 1.6um, 40°C, 0.4mL/min | 353 | 0 | 284 | 278 |
| | 0317 | Waters CORTECS UPLC C18 100x2.1mm, 1.6um, 40°C, 0.4mL/min | 141 | 0 | 74 | 77 |
| **Training datasets** | | | | | | |
| Publication - Pezzatti | 0179 | Phenomenex Kinetex C18 150x2.1mm, 1.7um, 30°C, 0.3mL/min | 307 | 316 | 78 | 90 |
| | 0180 | Phenomenex Kinetex C18 150x2.1mm, 1.7um, 30°C, 0.3mL/min | 273 | 310 | 69 | 77 |
| | 0181 | Phenomenex Kinetex C18 150x2.1mm, 1.7um, 30°C, 0.3mL/min | 214 | 466 | 86 | 89 |
| | 0182 | Phenomenex Kinetex C18 150x2.1mm, 1.7um, 30°C, 0.3mL/min | 164 | 159 | 41 | 41 |

| | | | | | | |
|---|---|---|---|---|---|---|
| Publication - Stoffel | 0187 | Merck Supelco Ascentis Express C18 100x2.1mm, 2um, 40°C, 0.3mL/min | 351 | 362 | 85 | 101 |
| | 0188 | Merck Supelco Ascentis Express C18 100x2.1mm, 2um, 40°C, 0.3mL/min | 330 | 308 | 75 | 84 |
| | 0189 | Merck Supelco Ascentis Express C18 100x2.1mm, 2um, 40°C, 0.2mL/min | 320 | 411 | 92 | 102 |
| | 0190 | Merck Supelco Ascentis Express C18 100x2.1mm, 2um, 40°C, 0.2mL/min | 303 | 315 | 69 | 89 |
| | 0191 | Merck Supelco Ascentis Express C18 100x2.1mm, 2um, 40°C, 0.25mL/min | 329 | 439 | 95 | 105 |
| | 0192 | Merck Supelco Ascentis Express C18 100x2.1mm, 2um, 40°C, 0.25mL/min | 309 | 298 | 69 | 88 |
| | 0193 | Merck Supelco Ascentis Express C18 100x2.1mm, 2um, 40°C, 0.35mL/min | 333 | 385 | 89 | 101 |
| | 0194 | Merck Supelco Ascentis Express C18 100x2.1mm, 2um, 40°C, 0.35mL/min | 313 | 309 | 72 | 86 |
| | 0195 | Merck Supelco Ascentis Express C18 100x2.1mm, 2um, 40°C, 0.4mL/min | 337 | 391 | 88 | 102 |
| | 0196 | Merck Supelco Ascentis Express C18 100x2.1mm, 2um, 40°C, 0.4mL/min | 308 | 328 | 70 | 91 |
| | 0197 | Merck Supelco Ascentis Express C18 100x2.1mm, 2um, 37.5°C, 0.3mL/min | 322 | 431 | 90 | 105 |
| | 0198 | Merck Supelco Ascentis Express C18 100x2.1mm, 2um, 37.5°C, 0.3mL/min | 303 | 335 | 73 | 93 |
| | 0199 | Merck Supelco Ascentis Express C18 100x2.1mm, 2um, 39°C, 0.3mL/min | 319 | 420 | 88 | 103 |
| | 0200 | Merck Supelco Ascentis Express C18 100x2.1mm, 2um, 39°C, 0.3mL/min | 298 | 340 | 70 | 92 |
| | 0201 | Merck Supelco Ascentis Express C18 100x2.1mm, 2um, 41°C, 0.3mL/min | 319 | 400 | 88 | 101 |
| | 0202 | Merck Supelco Ascentis Express C18 100x2.1mm, 2um, 41°C, 0.3mL/min | 297 | 350 | 69 | 93 |
| | 0203 | Merck Supelco Ascentis Express C18 100x2.1mm, 2um, 42.5°C, 0.3mL/min | 321 | 434 | 90 | 106 |
| | 0204 | Merck Supelco Ascentis Express C18 100x2.1mm, 2um, 42.5°C, 0.3mL/min | 299 | 334 | 69 | 93 |

| | | | | | | |
|---|---|---|---|---|---|---|
| Dataset - BGC | 0236 | Waters ACQUITY UPLC HSS T3 100x2.1mm, 1.8um, 30°C, 0.4mL/min | 446 | 1901 | 282 | 288 |
| | 0237 | Waters ACQUITY UPLC HSS T3 100x2.1mm, 1.8um, 30°C, 0.4mL/min | 318 | 1143 | 169 | 194 |
| | 0238 | Waters ACQUITY UPLC HSS T3 100x2.1mm, 1.8um, 40°C, 0.4mL/min | 461 | 1830 | 275 | 277 |
| | 0239 | Waters ACQUITY UPLC HSS T3 100x2.1mm, 1.8um, 40°C, 0.4mL/min | 322 | 1184 | 180 | 204 |
| | 0240 | Waters ACQUITY UPLC HSS T3 100x2.1mm, 1.8um, 50°C, 0.4mL/min | 449 | 1903 | 283 | 287 |
| | 0241 | Waters ACQUITY UPLC HSS T3 100x2.1mm, 1.8um, 50°C, 0.4mL/min | 294 | 1176 | 176 | 199 |
| | 0242 | Waters ACQUITY UPLC HSS T3 100x2.1mm, 1.8um, 40°C, 0.4mL/min | 319 | 325 | 79 | 83 |
| | 0243 | Waters ACQUITY UPLC HSS T3 100x2.1mm, 1.8um, 40°C, 0.4mL/min | 274 | 652 | 111 | 120 |
| | 0244 | Phenomenex Kinetex XB-C18 100x2.1mm, 1.7um, 30°C, 0.4mL/min | 451 | 397 | 101 | 112 |
| | 0245 | Phenomenex Kinetex XB-C18 100x2.1mm, 1.7um, 30°C, 0.4mL/min | 310 | 805 | 137 | 152 |
| | 0246 | Phenomenex Kinetex XB-C18 100x2.1mm, 1.7um, 40°C, 0.4mL/min | 475 | 387 | 104 | 113 |
| | 0247 | Phenomenex Kinetex XB-C18 100x2.1mm, 1.7um, 40°C, 0.4mL/min | 309 | 1354 | 193 | 211 |
| | 0248 | Phenomenex Kinetex XB-C18 100x2.1mm, 1.7um, 50°C, 0.4mL/min | 449 | 407 | 102 | 116 |
| | 0249 | Phenomenex Kinetex XB-C18 100x2.1mm, 1.7um, 50°C, 0.4mL/min | 305 | 800 | 136 | 151 |
| | 0250 | Phenomenex Kinetex XB-C18 100x2.1mm, 1.7um, 40°C, 0.4mL/min | 396 | 433 | 102 | 114 |
| | 0251 | Phenomenex Kinetex XB-C18 100x2.1mm, 1.7um, 40°C, 0.4mL/min | 277 | 616 | 101 | 114 |
| | 0252 | Waters ACQUITY UPLC BEH C18 100x2.1mm, 1.7um, 30°C, 0.4mL/min | 449 | 1929 | 290 | 288 |
| | 0253 | Waters ACQUITY UPLC BEH C18 100x2.1mm, 1.7um, 30°C, 0.4mL/min | 286 | 948 | 151 | 167 |

| | | | | | |
|---|---|---|---|---|---|
| 0254 | Waters ACQUITY UPLC BEH C18 100x2.1mm, 1.7um, 40°C, 0.4mL/min | 448 | 1927 | 286 | 283 |
| 0255 | Waters ACQUITY UPLC BEH C18 100x2.1mm, 1.7um, 40°C, 0.4mL/min | 75 | 368 | 57 | 46 |
| 0256 | Waters ACQUITY UPLC BEH C18 100x2.1mm, 1.7um, 50°C, 0.4mL/min | 448 | 2123 | 326 | 314 |
| 0257 | Waters ACQUITY UPLC BEH C18 100x2.1mm, 1.7um, 50°C, 0.4mL/min | 260 | 928 | 145 | 160 |
| 0258 | Waters ACQUITY UPLC BEH C18 100x2.1mm, 1.7um, 40°C, 0.4mL/min | 382 | 467 | 101 | 119 |
| 0259 | Waters ACQUITY UPLC BEH C18 100x2.1mm, 1.7um, 40°C, 0.4mL/min | 257 | 604 | 101 | 110 |
| 0326 | Restek Raptor Biphenyl 100x2.1mm, 2.7um, 30°C, 0.4mL/min | 21 | 0 | 5 | 2 |
| 0327 | Restek Raptor Biphenyl 100x2.1mm, 2.7um, 30°C, 0.4mL/min | 345 | 1193 | 182 | 201 |
| 0328 | Restek Raptor Biphenyl 100x2.1mm, 2.7um, 40°C, 0.4mL/min | 359 | 2002 | 278 | 280 |
| 0329 | Restek Raptor Biphenyl 100x2.1mm, 2.7um, 40°C, 0.4mL/min | 329 | 1259 | 193 | 211 |
| 0330 | Restek Raptor Biphenyl 100x2.1mm, 2.7um, 50°C, 0.4mL/min | 383 | 2249 | 318 | 320 |
| 0331 | Restek Raptor Biphenyl 100x2.1mm, 2.7um, 50°C, 0.4mL/min | 301 | 874 | 139 | 157 |
| 0332 | Restek Raptor Biphenyl 100x2.1mm, 2.7um, 40°C, 0.4mL/min | 381 | 2049 | 292 | 291 |
| 0333 | Restek Raptor Biphenyl 100x2.1mm, 2.7um, 40°C, 0.4mL/min | 171 | 437 | 70 | 79 |
| 0342 | Waters ACQUITY UPLC HSS C18 100x2.1mm, 1.8um, 30°C, 0.4mL/min | 24 | 0 | 4 | 2 |
| 0343 | Waters ACQUITY UPLC HSS C18 100x2.1mm, 1.8um, 30°C, 0.4mL/min | 309 | 1139 | 174 | 196 |
| 0344 | Waters ACQUITY UPLC HSS C18 100x2.1mm, 1.8um, 40°C, 0.4mL/min | 339 | 497 | 99 | 114 |
| 0345 | Waters ACQUITY UPLC HSS C18 100x2.1mm, 1.8um, 40°C, 0.4mL/min | 282 | 1206 | 176 | 199 |

| | | | | | | |
|---|---|---|---|---|---|---|
| | 0346 | Waters ACQUITY UPLC HSS C18 100x2.1mm, 1.8um, 50°C, 0.4mL/min | 356 | 673 | 131 | 137 |
| | 0347 | Waters ACQUITY UPLC HSS C18 100x2.1mm, 1.8um, 50°C, 0.4mL/min | 264 | 1250 | 184 | 202 |
| | 0348 | Waters ACQUITY UPLC HSS C18 100x2.1mm, 1.8um, 40°C, 0.4mL/min | 352 | 647 | 125 | 137 |
| | 0349 | Waters ACQUITY UPLC HSS C18 100x2.1mm, 1.8um, 40°C, 0.4mL/min | 174 | 444 | 75 | 78 |
| Dataset - Rutz | 0264 | Waters ACQUITY UPLC BEH C18 50x2.1mm, 1.7um, 40°C, 0.6mL/min | 187 | 0 | 24 | 24 |
| MZmine | 0390 | Waters ACQUITY UPLC BEH C18 50x2.1mm, 1.7um, 40°C, 0.6mL/min | 2889 | 0 | 309 | 369 |
| | 0391 | Waters ACQUITY UPLC BEH C18 50x2.1mm, 1.7um, 40°C, 0.6mL/min | 5672 | 286 | 727 | 737 |

Table S2: Datasets evaluation summary. Here $N$ stands for the number of datapoints in the subset.

| Evaluation datasets | | | | | | | | | | | | |
|---|---|---|---|---|---|---|---|---|---|---|---|---|
| RepoRT ID | $Mol_{train}$ | | | | $Mol_{val}$ | | | | $Mol_{test}$ | | | |
| | $N$ | MAE | MRE | $R^2$ | $N$ | MAE | MRE | $R^2$ | $N$ | MAE | MRE | $R^2$ |
| 0311 | 303 | 0.26 | 0.13 | 0.96 | 38 | 0.42 | 0.18 | 0.90 | 50 | 0.59 | 0.23 | 0.78 |
| 0312 | 349 | 0.23 | 0.09 | 0.98 | 48 | 0.48 | 0.25 | 0.83 | 46 | 0.67 | 0.22 | 0.79 |
| 0313 | 294 | 0.25 | 0.13 | 0.95 | 42 | 0.42 | 0.22 | 0.89 | 50 | 0.63 | 0.27 | 0.76 |
| 0314 | 365 | 0.26 | 0.10 | 0.97 | 48 | 0.49 | 0.25 | 0.80 | 45 | 0.70 | 0.22 | 0.77 |
| 0315 | 268 | 0.22 | 0.12 | 0.98 | 34 | 0.45 | 0.24 | 0.86 | 41 | 0.73 | 0.27 | 0.67 |
| 0316 | 355 | 0.23 | 0.09 | 0.96 | 47 | 0.49 | 0.25 | 0.80 | 44 | 0.59 | 0.20 | 0.83 |
| 0317 | 144 | 0.22 | 0.09 | 0.98 | 19 | 0.59 | 0.18 | 0.61 | 9 | 0.41 | 0.13 | 0.80 |
| 0389 | 65 | 1.08 | 0.30 | 0.92 | 9 | 1.13 | 0.46 | 0.92 | 9 | 1.43 | 0.15 | 0.87 |
| Training datasets | | | | | | | | | | | | |
| RepoRT ID | $Mol_{train}$ | | | | $Mol_{val}$ | | | | $Mol_{test}$ | | | |
| | $N$ | MAE | MRE | $R^2$ | $N$ | MAE | MRE | $R^2$ | $N$ | MAE | MRE | $R^2$ |
| 0179 | 319 | 0.40 | 0.17 | 0.96 | 41 | 0.39 | 0.18 | 0.93 | 41 | 0.40 | 0.20 | 0.95 |
| 0180 | 285 | 0.37 | 0.15 | 0.90 | 38 | 0.38 | 0.17 | 0.94 | 38 | 0.42 | 0.20 | 0.94 |
| 0181 | 219 | 0.41 | 0.17 | 0.95 | 23 | 0.40 | 0.21 | 0.87 | 25 | 0.36 | 0.18 | 0.89 |
| 0182 | 166 | 0.41 | 0.16 | 0.96 | 20 | 0.46 | 0.18 | 0.95 | 20 | 0.37 | 0.18 | 0.92 |
| 0187 | 364 | 0.24 | 0.09 | 0.98 | 46 | 0.51 | 0.20 | 0.75 | 48 | 0.34 | 0.15 | 0.91 |
| 0188 | 344 | 0.22 | 0.07 | 0.98 | 42 | 0.27 | 0.16 | 0.93 | 44 | 0.27 | 0.10 | 0.91 |
| 0189 | 333 | 0.18 | 0.06 | 0.99 | 41 | 0.59 | 0.16 | 0.77 | 42 | 0.31 | 0.13 | 0.89 |

| | | | | | | | | | | | | |
|---|---|---|---|---|---|---|---|---|---|---|---|---|
| 0190 | 314 | 0.16 | 0.05 | 0.99 | 39 | 0.36 | 0.15 | 0.92 | 39 | 0.29 | 0.08 | 0.92 |
| 0191 | 342 | 0.17 | 0.06 | 0.99 | 43 | 0.44 | 0.17 | 0.85 | 46 | 0.28 | 0.14 | 0.90 |
| 0192 | 320 | 0.15 | 0.05 | 0.99 | 41 | 0.35 | 0.16 | 0.91 | 39 | 0.26 | 0.08 | 0.92 |
| 0193 | 345 | 0.21 | 0.08 | 0.98 | 43 | 0.48 | 0.20 | 0.84 | 44 | 0.23 | 0.15 | 0.92 |
| 0194 | 324 | 0.20 | 0.08 | 0.99 | 40 | 0.32 | 0.20 | 0.91 | 42 | 0.21 | 0.09 | 0.94 |
| 0195 | 350 | 0.24 | 0.09 | 0.98 | 43 | 0.44 | 0.22 | 0.82 | 45 | 0.22 | 0.15 | 0.93 |
| 0196 | 321 | 0.20 | 0.09 | 0.99 | 41 | 0.31 | 0.22 | 0.91 | 40 | 0.21 | 0.10 | 0.94 |
| 0197 | 332 | 0.19 | 0.07 | 0.98 | 42 | 0.44 | 0.19 | 0.84 | 40 | 0.31 | 0.16 | 0.89 |
| 0198 | 313 | 0.16 | 0.06 | 0.99 | 39 | 0.34 | 0.18 | 0.91 | 37 | 0.26 | 0.08 | 0.93 |
| 0199 | 330 | 0.18 | 0.07 | 0.98 | 40 | 0.46 | 0.19 | 0.84 | 41 | 0.30 | 0.16 | 0.89 |
| 0200 | 310 | 0.15 | 0.06 | 0.99 | 39 | 0.33 | 0.18 | 0.91 | 38 | 0.25 | 0.08 | 0.93 |
| 0201 | 332 | 0.23 | 0.06 | 0.91 | 41 | 0.40 | 0.19 | 0.85 | 40 | 0.26 | 0.15 | 0.91 |
| 0202 | 308 | 0.15 | 0.07 | 0.99 | 38 | 0.33 | 0.19 | 0.91 | 38 | 0.24 | 0.08 | 0.93 |
| 0203 | 334 | 0.17 | 0.06 | 0.98 | 42 | 0.47 | 0.20 | 0.83 | 40 | 0.29 | 0.16 | 0.89 |
| 0204 | 310 | 0.14 | 0.06 | 0.99 | 38 | 0.33 | 0.19 | 0.91 | 38 | 0.24 | 0.08 | 0.92 |
| 0236 | 450 | 0.20 | 0.07 | 0.97 | 55 | 0.42 | 0.16 | 0.87 | 56 | 0.47 | 0.15 | 0.91 |
| 0237 | 327 | 0.20 | 0.08 | 0.97 | 39 | 0.39 | 0.13 | 0.89 | 40 | 0.38 | 0.16 | 0.92 |
| 0238 | 465 | 0.20 | 0.07 | 0.98 | 57 | 0.40 | 0.15 | 0.88 | 58 | 0.49 | 0.15 | 0.91 |
| 0239 | 331 | 0.17 | 0.07 | 0.98 | 45 | 0.40 | 0.17 | 0.88 | 46 | 0.39 | 0.16 | 0.91 |
| 0240 | 451 | 0.18 | 0.07 | 0.98 | 57 | 0.42 | 0.16 | 0.87 | 52 | 0.47 | 0.15 | 0.90 |
| 0241 | 300 | 0.15 | 0.07 | 0.99 | 42 | 0.37 | 0.13 | 0.90 | 38 | 0.39 | 0.17 | 0.89 |
| 0242 | 322 | 0.22 | 0.07 | 0.96 | 45 | 0.46 | 0.15 | 0.84 | 42 | 0.55 | 0.21 | 0.87 |
| 0243 | 282 | 0.21 | 0.08 | 0.95 | 35 | 0.46 | 0.17 | 0.85 | 34 | 0.41 | 0.15 | 0.88 |
| 0244 | 455 | 0.21 | 0.10 | 0.95 | 54 | 0.39 | 0.19 | 0.91 | 58 | 0.62 | 0.24 | 0.78 |
| 0245 | 320 | 0.17 | 0.09 | 0.97 | 38 | 0.38 | 0.15 | 0.90 | 40 | 0.40 | 0.22 | 0.86 |
| 0246 | 478 | 0.20 | 0.09 | 0.96 | 56 | 0.36 | 0.18 | 0.92 | 59 | 0.58 | 0.22 | 0.82 |
| 0247 | 318 | 0.17 | 0.08 | 0.98 | 40 | 0.40 | 0.21 | 0.89 | 41 | 0.41 | 0.21 | 0.85 |
| 0248 | 452 | 0.19 | 0.08 | 0.95 | 57 | 0.46 | 0.20 | 0.86 | 57 | 0.62 | 0.22 | 0.76 |
| 0249 | 312 | 0.15 | 0.08 | 0.98 | 40 | 0.37 | 0.20 | 0.89 | 39 | 0.39 | 0.19 | 0.85 |
| 0250 | 398 | 0.23 | 0.09 | 0.94 | 50 | 0.39 | 0.13 | 0.87 | 45 | 0.63 | 0.22 | 0.76 |
| 0251 | 284 | 0.19 | 0.08 | 0.97 | 33 | 0.36 | 0.14 | 0.90 | 34 | 0.42 | 0.18 | 0.85 |
| 0252 | 454 | 0.18 | 0.08 | 0.95 | 57 | 0.47 | 0.19 | 0.85 | 58 | 0.39 | 0.13 | 0.94 |
| 0253 | 295 | 0.16 | 0.09 | 0.98 | 36 | 0.34 | 0.14 | 0.93 | 35 | 0.37 | 0.18 | 0.92 |
| 0254 | 452 | 0.20 | 0.08 | 0.94 | 55 | 0.43 | 0.19 | 0.85 | 57 | 0.41 | 0.14 | 0.93 |
| 0255 | 75 | 0.11 | 0.03 | 0.98 | 6 | 0.68 | 0.24 | -0.02 | 6 | 0.33 | 0.15 | 0.89 |
| 0256 | 452 | 0.17 | 0.06 | 0.96 | 55 | 0.47 | 0.15 | 0.84 | 59 | 0.37 | 0.11 | 0.94 |
| 0257 | 268 | 0.14 | 0.08 | 0.99 | 32 | 0.36 | 0.16 | 0.88 | 33 | 0.36 | 0.18 | 0.92 |
| 0258 | 387 | 0.20 | 0.07 | 0.97 | 47 | 0.42 | 0.13 | 0.84 | 40 | 0.35 | 0.11 | 0.93 |
| 0259 | 272 | 0.13 | 0.06 | 0.99 | 33 | 0.37 | 0.20 | 0.91 | 31 | 0.38 | 0.15 | 0.91 |
| 0264 | 187 | 0.48 | 0.42 | 0.84 | 24 | 0.60 | 0.27 | 0.88 | 24 | 0.51 | 0.40 | 0.79 |
| 0310 | 19 | 1.00 | 0.29 | 0.18 | 2 | 0.26 | 0.06 | -inf | 2 | 0.53 | 0.10 | -41.17 |
| 0326 | 22 | 0.85 | 0.32 | 0.50 | 5 | 0.57 | 0.27 | 0.70 | 5 | 0.77 | 0.46 | 0.67 |
| 0327 | 354 | 0.18 | 0.09 | 0.98 | 43 | 0.31 | 0.14 | 0.90 | 51 | 0.39 | 0.17 | 0.89 |
| 0328 | 359 | 0.16 | 0.07 | 0.98 | 49 | 0.36 | 0.17 | 0.89 | 47 | 0.36 | 0.14 | 0.94 |
| 0329 | 337 | 0.17 | 0.09 | 0.98 | 43 | 0.33 | 0.15 | 0.89 | 47 | 0.42 | 0.19 | 0.88 |
| 0330 | 383 | 0.16 | 0.07 | 0.98 | 51 | 0.35 | 0.16 | 0.89 | 46 | 0.36 | 0.14 | 0.93 |
| 0331 | 309 | 0.16 | 0.09 | 0.98 | 41 | 0.32 | 0.14 | 0.91 | 45 | 0.39 | 0.20 | 0.88 |
| 0332 | 381 | 0.19 | 0.08 | 0.98 | 50 | 0.44 | 0.18 | 0.82 | 45 | 0.38 | 0.15 | 0.92 |
| 0333 | 173 | 0.18 | 0.08 | 0.95 | 24 | 0.32 | 0.12 | 0.85 | 14 | 0.40 | 0.20 | 0.83 |
| 0342 | 25 | 0.88 | 0.32 | 0.64 | 4 | 0.86 | 0.36 | 0.39 | 5 | 0.79 | 0.48 | 0.72 |
| 0343 | 317 | 0.18 | 0.09 | 0.96 | 37 | 0.42 | 0.20 | 0.85 | 45 | 0.55 | 0.19 | 0.84 |
| 0344 | 340 | 0.12 | 0.07 | 0.99 | 40 | 0.34 | 0.18 | 0.92 | 44 | 0.54 | 0.16 | 0.88 |
| 0345 | 291 | 0.15 | 0.08 | 0.97 | 37 | 0.39 | 0.19 | 0.88 | 50 | 0.49 | 0.18 | 0.87 |
| 0346 | 357 | 0.12 | 0.06 | 0.99 | 49 | 0.42 | 0.19 | 0.85 | 45 | 0.53 | 0.16 | 0.88 |
| 0347 | 274 | 0.12 | 0.07 | 0.98 | 36 | 0.38 | 0.19 | 0.87 | 40 | 0.40 | 0.16 | 0.89 |
| 0348 | 353 | 0.14 | 0.07 | 0.98 | 42 | 0.38 | 0.14 | 0.88 | 46 | 0.44 | 0.14 | 0.91 |

| | | | | | | | | | | | | |
|---|---|---|---|---|---|---|---|---|---|---|---|---|
| 0349 | 177 | 0.15 | 0.07 | 0.94 | 21 | 0.39 | 0.12 | 0.84 | 16 | 0.26 | 0.12 | 0.94 |
| 0390 | 5895 | 0.32 | 0.11 | 0.87 | 719 | 0.42 | 0.13 | 0.82 | 719 | 0.40 | 0.13 | 0.81 |
| 0391 | 7810 | 0.34 | 0.12 | 0.85 | 937 | 0.41 | 0.15 | 0.77 | 936 | 0.50 | 0.16 | 0.69 |

### References


[1] Daniel Servén and Charlie Brummitt. pygam: Generalized additive models in python, March 2018.

[2] Jan Stanstrup, Steffen Neumann, and Urska Vrhovsek. Predret: prediction of retention time by direct mapping between multiple chromatographic systems. *Analytical chemistry*, 87(18):9421–9428, 2015.

[3] F. Pedregosa, G. Varoquaux, A. Gramfort, V. Michel, B. Thirion, O. Grisel, M. Blondel, P. Prettenhofer, R. Weiss, V. Dubourg, J. Vanderplas, A. Passos, D. Cournapeau, M. Brucher, M. Perrot, and E. Duchesnay. Scikit-learn: Machine learning in Python. *Journal of Machine Learning Research*, 12:2825–2830, 2011.

[4] LR Snyder, JW Dolan, and PW Carr. The hydrophobic-subtraction model of reversed-phase column selectivity. *Journal of Chromatography A*, 1060(1-2):77–116, 2004.

[5] Qiong Yang, Hongchao Ji, Hongmei Lu, and Zhimin Zhang. Prediction of liquid chromatographic retention time with graph neural networks to assist in small molecule identification. *Analytical Chemistry*, 93(4):2200–2206, 2021.
